# From Adipose to Limbus: Deciphering the Paracrine Effects of MSC Secretomes on Oxidative Stress-Induced RPE Dysfunction

**DOI:** 10.64898/2026.03.24.707782

**Authors:** Ayşe Dilara Aydemir, Zehra Canbulat, Murat Hasanreisoğlu

**Affiliations:** Koç University Research Center for Translational Medicine (KUTTAM), Koç University, 34450, Istanbul, Türkiye; Department of Ophthalmology, Koç University School of Medicine, Istanbul, Türkiye

## Abstract

This study investigates the therapeutic potential of secretomes derived from Adipose-derived Mesenchymal Stem Cells (ADMSC-CM) and Limbal-derived Mesenchymal Stem Cells (LMSC-CM) against oxidative stress-induced damage in Retinal Pigment Epithelium (RPE-1) cells. RPE dysfunction, often triggered by oxidative stress, is a hallmark of various retinal degenerations. Here, we induced RPE-1 injury using H_2_O_2_ and evaluated the restorative effects of both MSC-conditioned media (CM). Our results demonstrated that both ADMSC-CM and LMSC-CM significantly enhanced cell viability and successfully reversed H_2_O_2_-induced G2/M phase cell cycle arrest. While oxidative stress triggered a pro-inflammatory response characterized by elevated IL−1β, IL−6, and IL−10 expression, MSC-CM treatment, particularly ADMSC-CM, effectively modulated these levels and suppressed the p38 MAPK signaling pathway. Furthermore, MSC-CM reduced the Bax/Bcl-2 ratio, indicating an anti-apoptotic effect, and appeared to stabilize autophagic flux. To investigate the impact of oxidative-stress induced alterations in retinal pigment epithelial cells on angiogenesis, the effects of RPE-derived secreted factors on endothelial cell function were evaluated. Crucially, in terms of safety and secondary complications, neither secretome exhibited pro-angiogenic tendencies; instead, they significantly inhibited HUVEC migration and invasion compared to the H_2_O_2_ damaged group. These findings suggest that both ADMSC and LMSC secretomes provide a potent multi-targeted therapeutic effect, making them promising candidates for cell-free therapies in retinal diseases.

## INTRODUCTION

The retinal pigment epithelium (RPE) serves as a critical guardian of retinal homeostasis, managing everything from nutrient transport to the phagocytosis of photoreceptor outer segments (Strauss, 2005). However, its continuous exposure to light and elevated metabolic activity render it a principal target for oxidative stress. Over time, the accumulation of reactive oxygen species (ROS) results in retinal pigment epithelium (RPE) senescence, inflammation, and ultimately, the permanent vision loss associated with Age-Related Macular Degeneration (AMD) (Goldfischer et al., 1966; Lim et al., 2012; Marmorstein, 2001).

Conventional cell-based therapies have demonstrated potential; yet, concerns about immunological rejection and surgical integration have redirected attention to the secretome, a potent mixture of growth factors and cytokines released by Mesenchymal Stem Cells (MSCs) (Martinez-Rodriguez et al., 2025). While adipose-derived MSCs (ADMSCs) are well-known for their robust immunomodulatory properties (Dam et al., 2026; El-Sayed et al., 2019), limbal-derived MSCs (LMSCs) offer a unique, tissue-specific alternative due to their proximity to ocular structures (Nili et al., 2019; Tavakkoli et al., 2022). However, a direct comparison of their therapeutic efficacy in the context of RPE recovery remains largely unexplored.

This work aimed to address this gap by assessing the impact of ADMSC and LMSC secretomes on the recovery of H_2_O_2_-treated RPE cells. In addition to basic cell survival, we investigated the mechanisms involved: how these secretomes influence the cell cycle, their potential to mitigate the inflammatory senescence-associated secretory phenotype (SASP) through the p38 MAPK pathway, and their function in regulating autophagy and death. Furthermore, given the risk of neovascularization in retinal pathologies, we investigated the impact of these treatments on angiogenic processes. By providing a comprehensive comparison of these two MSC sources, this research seeks to identify the most effective and safe cell-free strategy for treating oxidative damage in the retina.

## MATERIALS AND METHODS

### Ethics Statements

The protocols for tissue collection and primary cell isolation in this study were strictly conducted in accordance with the 1964 Declaration of Helsinki and its subsequent amendments. Ethical approval was granted by the Koç University Clinical Research Ethics Committee (Institutional Review Board) under the following protocols: 2020.447.IRB2.121 for adipose tissue and 2019.090.IRB2.027 for limbal tissue. Human adipose tissues were obtained form healthy donors undergoing elective liposuction surgery at Koç University Hospital. All participants provided written informed consent after being fully briefed on the nature, scope, and potential implications of the research. Limbal tissues were isolated from cadaveric corneascleral rims provided by the Ministry of Health Cornea Bank, Turkey. These tissues remained after the central corneal buttons were utilized for transplantation.

### Research Design and Approach

Given that MSCs derived from different tissues exhibit varying effects, this study investigated the effects of CM from adipose tissue and limbal-derived stem cells on RPE cells subjected to H_2_O_2_-induced stress. The study examined the impact of the CM on cell viability, cell cycle arrest, inflammation, and autophagy in RPE cells. Additionally, the angiogenic effects were assessed using human umbilical vein endothelial cells (HUVECs) by evaluating cell viability, migration, and invasion, along with VEGF release determined by ELISA (Fig. 1).

**Figure 1.**
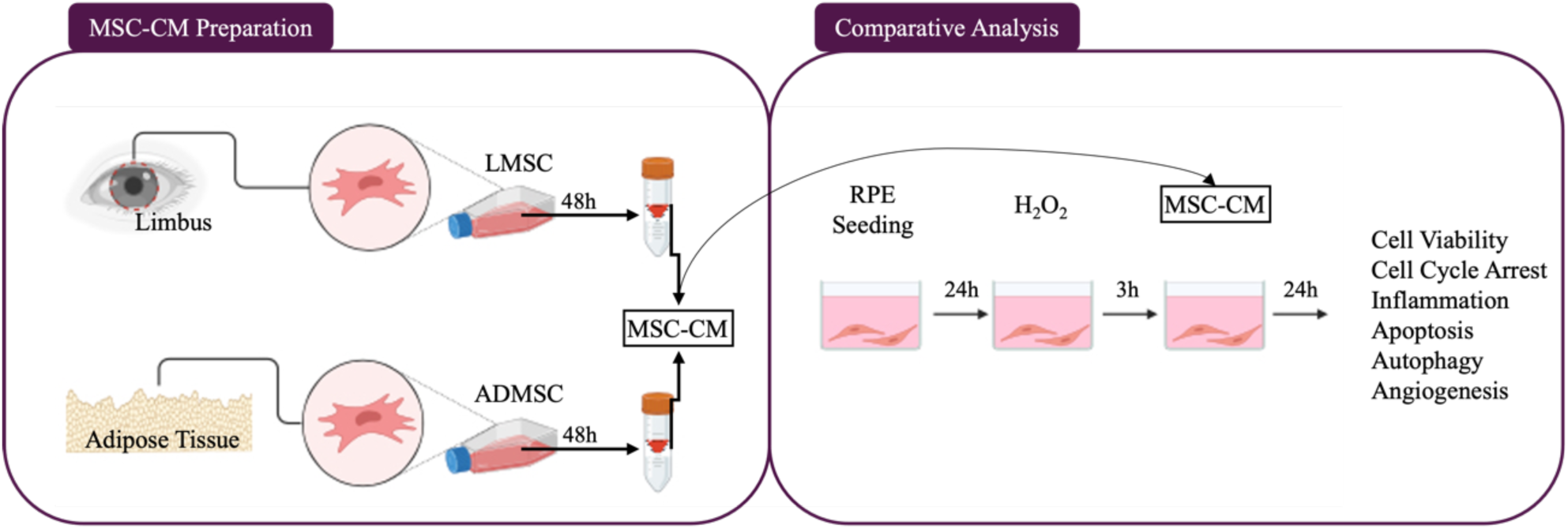
The Research Design. Comparative effect of MSC-CM therapy was assessed by using the concentrated mesenchymal stem cell-conditioned medium derived from adipose tissue and limbal tissue (Created with Biorender.com).

### Isolation and Characterization of Mesenchymal Stem Cell

#### Isolation and Characterization of Adipose-Derived Mesenchymal Stem Cell

ADMSCs were isolated and characterized as previously described (Gozel et al., 2024). Adipose tissues from liposuction were washed with DPBS (Biowest, Nuaille, France) and enzymatically digested with 0.075% collagenase I at 37°C. The suspension was neutralized with DMEM-LG (10% FBS, 1% P/S), centrifuged at 800 g for 10 minutes, and filtered through a 70 µm mesh. Cells were then seeded into T25 flasks and cultured at 37°C with 5% CO_2_. Upon 70–80% confluency, cells were passaged using 0.25% Trypsin-EDTA. Characterization was performed at P3 via lineage-specific staining (Alizarin Red, Alcian Blue, and Oil Red O) to confirm tri-lineage differentiation. MSCs at P5 were utilized for conditioned media (CM) preparation.

#### Isolation and Characterization of Limbal Mesenchymal Stem Cell

LMSCs were isolated and characterized following the established protocol previously reported by our group (Gulzar et al., 2022). Briefly, cadaveric corneoscleral rims were transported in Optisol medium and washed with PBS containing 1% P/S. After the removal of the epithelial layer, the tissue was divided into four equal explants and placed onto Matrigel-coated 12-well plates. The explants were initially cultured in 300 µl of complete medium, with an additional 200 µl added after 24 hours to promote cell attachment. The culture medium was refreshed every three days. Upon observing significant cell migration from the tissue (P0), cells were expanded into T25 flasks (P1) using 0.25% Trypsin-EDTA once 70–80% confluency was reached. Consistent with the ADMSC protocol, LMSCs at P3 were characterized via Alizarin Red, Alcian Blue, and Oil Red O staining to confirm their tri-lineage differentiation potential. Cells at P5 were utilized for the preparation of conditioned media (CM).

#### Collection and Preparation of Mesenchymal Stem Cell-Conditioned Medium

ADMSCs and LMSCs were grown until they attained 80% confluency. Prior to CM collection, the cells were grown in serum-free DMEM-LG for 48 hours. Following culturing in serum-free DMEM-LG, the culture media was collected into a 50 mL centrifuge tube and centrifuged at 1500xg for 10 minutes to remove cell debris. Centrifugal filter units with a molecular mass cut-off of 100 kDa (Amicon® Ultra-15, Millipore) were used to concentrate the centrifuged supernatant 50-fold. The supernatant was centrifuged for 40 minutes at 4000 g at room temperature. The concentrated CM was collected into 1.5 mL Eppendorf tubes (Eppendorf, HK) and stored at-80°C until use. The CM used in each experimental setup was isolated using MSCs isolated from different tissues. The Pierce™ BCA Protein Assay Kit (Thermo Fisher Scientific, MA, USA) was used to determine the protein content.

#### Cell Culture and Experimental Setup

The human RPE cell line RPE-1 were cultured at 37°C, 5% CO2 in Dulbecco’s Modified Eagle Medium/Nutrient Mixture F-12 (DMEM-F12) with 10% fetal bovine serum. RPE-1 were used in passage 12-17. Cells were passaged when they reached 90% confluence through trypsinization with 0.25% trypsin-EDTA and seeded into suitable culture plates for the next experiments. Freshly prepared solutions of H_2_O_2_, MSC-CM, and chloroquine (CQ) were added to the cell culture medium.

#### Determination of Cell Viability

The 3-(4,5-dimethylthiazol-2-yl)-2,5-diphenyltetrazolium bromide] cell viability assay was used to assess the cell viability of RPE-1 cells. In 96-well plates, 1×10^4^ cells/well were seeded, which were then treated with H_2_O_2_ in the presence or absence of MSC-conditioned media. Following treatment, MTT (0.5 mg/ml) was added to each well, and then incubated for a further 2h at 37°C prior to the addition of dimethyl sulfoxide. Absorbance was measured at 570 nm according to the manufacturer’s protocol using Synergy H1 Microplate Reader (BioTek). The percentage of cell viability was then determined by comparing it to the control group. All experiments were performed in quintuplicate.

#### Cell Cycle Distribution Analysis

The adherent cells were washed with PBS, collected using a trypsin-EDTA solution, and then suspended in PBS at a concentration of 1×10^8^ cells/ml. Afterwards, FxCycle^TM^PI/ RNase Staining solution (Thermo Fisher Scientific) were added to each sample tube. Following incubation for 10min at room temperature, the samples were analyzed using an Attune Nxt Acoustic Focusing Flow Cytometer system (Thermo Fisher Scientific). Cell cycle distribution analysis was performed by FlowJo v10.10.0 (BD Biosciences, USA).

#### Evaluation of Changes in Protein Expression

Cells were washed 2 times with PBS and the cell pellet for protein isolation were incubated for 30 minutes in ice with RIPA buffer (Thermo Fisher Scientific, Waltham, MA, USA). Following incubation, the samples were centrifuged at 16500 g for 15 minutes at 4°C and the the supernatants were transferred to new tubes. Protein concentrations were determined with the Bradford protein assay kit (Thermo Fisher Scientific, MA, USA). Isolated amounts of protein (10 µg for all samples) were loaded onto a 4-12% polyacrylamide gel and after run-in the proteins in the gel will be transferred to the Immuno-Blot® PVDF membrane (Bio-Rad, Hercules, CA, USA). The membrane was treated with 5% skim milk in TBS-T for 60 minutes to avoid non-specific binding. The membranes with the primary antibodies were incubated at 4°C overnight. Table 1 lists the detail of antibodies used for western blot. Following washing with TBS-T, membranes were incubated with horseradish peroxidase-conjugated anti-rabbit and anti-mouse IgG secondary antibodies for 60 minutes at room temperature. After treatment, membranes were washed with TBS-T and protein-antibody complexes were visualized with PierceTM ECL Western Blotting Substrate. Protein bands on the membrane were observed with the Bio-Rad ChemiDocTM MP Imaging System (Hercules, CA, USA).

**Table 1:** List of antibodies used in western blotting.

| Antibody Name | Company/Cat No | Dilution |
| --- | --- | --- |
| anti-p38 | Abcam/ab170099 | 1:1000 |
| anti-phospho-p38 | Abcam/ab45381 | 1:1000 |
| anti-Bax | Abcam/ab216494 | 1:1000 |
| anti-Bcl-2 | Abcam/ab59348 | 1:1000 |
| anti-LC3B | Novus Biologicals/NB00-1384 | 1:1000 |
| anti-p62 | Cell Signaling Technology/cs-5114 | 1:1000 |
| anti-vinculin | Abcam/ab130007 | 1:1000 |

### Evaluation of Angiogenic Function

#### Quantification of VEGF release

VEGF-A in the lysates was quantified using an ELISA MAX™ Deluxe Set Human VEGF (Biolegend), following the manufacturer’s instructions.

#### Determination of HUVEC Viability

The 3-(4,5-dimethylthiazol-2-yl)-2,5-diphenyltetrazolium bromide] cell viability assay was used to assess the cell viability of Human Umbilical Vein Endothelial Cells (HUVECs). Following the administration of RPE-1 treatment, samples of control, H_2_O_2_, and H_2_O_2_+MSC-CM supernatants were obtained and subsequently applied to endothelial cells to further evaluate the effects of the secretome on angiogenesis. In 96-well plates, 1.5×10^4^ cells/well were seeded. Following treatment, MTT Assay was performed.

#### Effect on Endothelial Cell Migration

A 24-well plate was used to seed the HUVECs. A sterile 200-µL pipette tip was used to make a straight incision in the middle of the monolayer once the cells reached 80% confluence. After two rounds of PBS washing, the wells were cultured for 24 hours with a 1:1 mixture of HUVEC and RPE-1 treatment media to eliminate any dead cells. Using an inverted microscope (Nikon, Tokyo, Japan), the wounds were imaged using phase contrast microscopy. Wound healing rate was calculated by subtracting the final wound size from the initial wound size and dividing the result by the initial wound size. The Fiji software was used to determine the initial and final wound sizes. A total of three separate trials were conducted.

#### Effect on Endothelial Cell Invasion

To evaluate the effect of supernatant on the invasion of HUVECs, a Transwell system was employed. HUVECs were seeded in the upper compartment of the Transwell inserts at a density of 2.3 × 10⁴ cells per well. The cells were maintained in an incubator set at 37°C with 5% CO_2_ The seeding medium contained 1% FBS. In the lower compartment of the Transwell inserts, the supernatant was added. As a positive control, medium containing 10% FBS was used. The cells were allowed to invade towards the lower compartment for a duration of 6 hours. After the 6-hour incubation period, non-invaded cells in the upper compartment were gently wiped off. The adherent cells were then fixed using 2% PFA for 10 minutes. Following fixation, the adherent cells were stained with crystal violet for 5 minutes. After staining, the inserts were washed with PBS and left to air dry. Images of the stained cells were captured using an inverted microscope, and the cells were counted manually.

### Data Collection and Statistical Analysis

All data are from at least three independent experiments and are expressed as mean ± standard error of the mean (SEM). Statistical evaluation of the data was performed using One-way ANOVA and unpaired t-tests to compare the mean of parametric distributed metric data between than two groups and more two groups, respectively, and Mann–Whitney U-test for non-parametric distributed metric data. Statistical significance was defined as P < 0.05. The statistical program GraphPad Prism 8 (GraphPad Software, San Diego, CA, USA) was used to do the statistical analysis

## RESULTS

### MSC-CM elevated RPE cell viability without affecting their ability to proliferate

Dose of MSC-CM were determined by observing its effect on the viability of RPE-1 cells. Cell viability was evaluated by MTT assay method. It was determined that MSC-CM initially applied to control cells did not have an adverse effect at any dose. On the contrary, a statistically significant increase in cell viability was observed in 200 µg ADMSC-CM and 50 µg LMSC-CM treated group compared to the control group. Interestingly, when 200 µg of LMSC-CM was administered, cell viability significantly decreased compared to the 200 µg ADMSC-CM and 50 µg LMSC-CM groups. However, this decrease was not below the % cell viability of the control group (Fig.2A). Considering these findings, the MSC-CM dose and application time was determined based on the survival rates H_2_O_2_-treated RPE-1 cells. Following 3-hour application of 400 µM H_2_O_2_ a decrease in cell survival was observed, and this decrease was elevated at 24 after the application was terminated in the basal group (Fig. 2B). With application of 200 µg ADMSC-CM resulted in significantly increased cell survival relative to the H_2_O_2_group (p< 0.001; Fig. 2B). On the other hand, LMSC-CM application increased survival significantly compared to the H_2_O_2_ group at both concentrations (p< 0.05; Fig. 2B). Finally, no significant differences were observed between both CM applications at both concentrations after H_2_O_2_ treatment (Fig. 2B). As a result of these results, subsequent assays were performed with a 24h treatment application using 200µg MSC-CM on H_2_O_2_ treated RPE-1 cells. Then, the proliferation effect of MSC-CM on H_2_O_2_-treated RPE-1 cells was observed by Ki67 staining method (Fig. 2C). H_2_O_2_ treatment did not negatively affect the proliferative abilities of RPE-1 cells. As intended, the 200 µg MSC-CM treatment groups did not significantly enhance the proliferation abilities of H_2_O_2_-treated RPE-1 cells (Fig. 2D).

**Figure 2.**
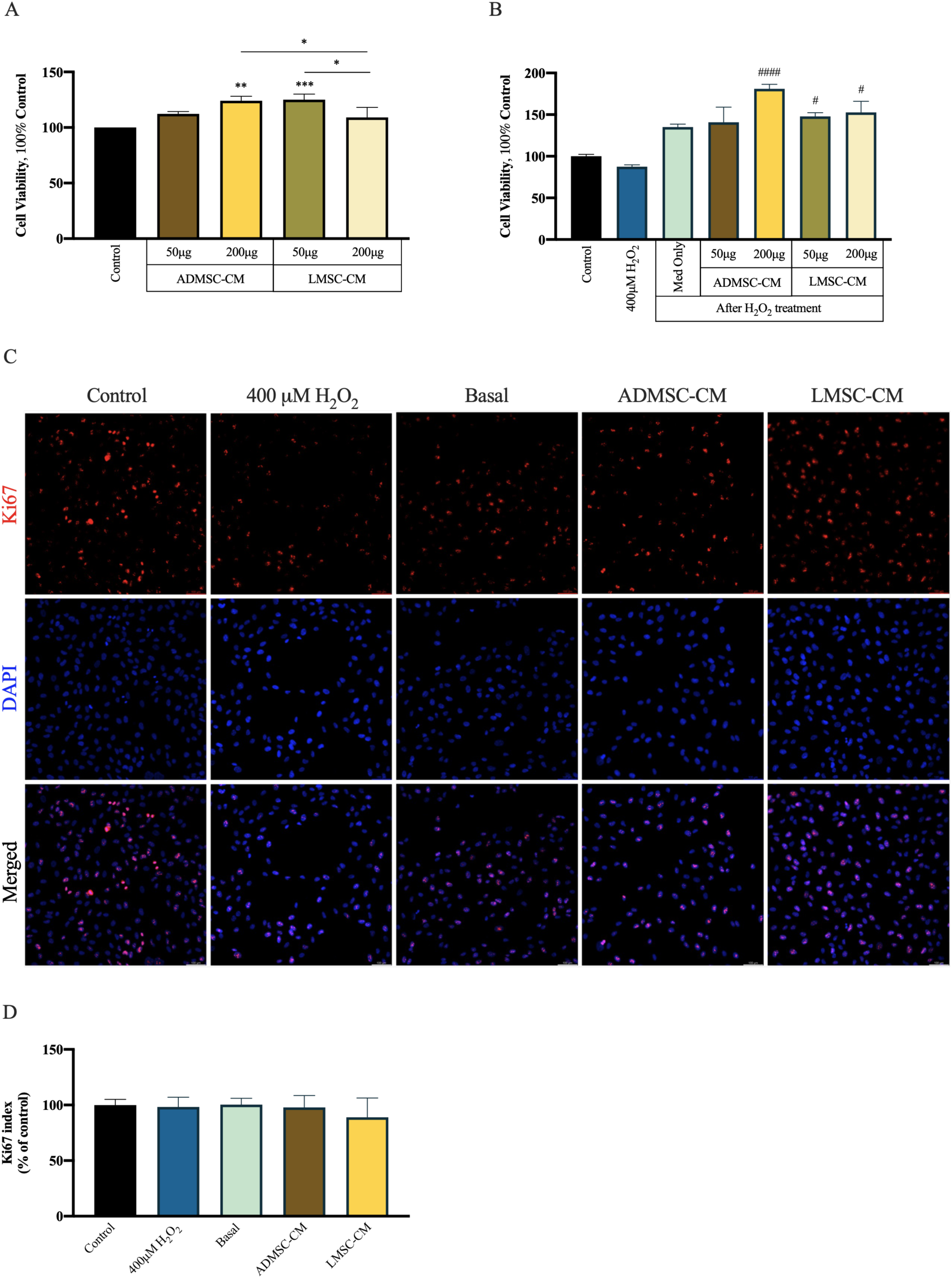
MSC-CM improved H₂O₂-induced cytotoxicity. A) RPE cell viability was assessed using MTT assay after treatment with adipose and limbal tissue-derived MSC-CM for 24 hours at indicated doses. The results are expressed as percentage of control. B) Cell viability was measured by MTT assay in H₂O₂-treated cells at 24 h after different doses of both MSC-CM treatments. The results are expressed as percentage of control. C) The proliferation effect of ADMSC-CM and LMSC-CM on H₂O₂-treated cells at 24h time-course was showed by Ki67 staining. Scale bar: 100 µm D) Ki67 index in RPE cells. Statistical analysis was performed by One-way ANOVA test. Data are expressed as mean ± SD. * compared to control group, # compared to H₂O₂ group; for all comparisons * p<0.05, ** p<0.01, *** p<0.001, **** p<0.0001.

### MSC-CM regulated impaired G2/M arrest

The cell cycle distribution profile was evaluated in order to investigate the mechanism underlying the MSC-CM-induced viability effect in H_2_O_2_-treated RPE-1 cells. Representative images of cell cycle analysis results summarized in Figure 3. The results showed that G2/M phase cell cycle arrest occurred after H_2_O_2_ treatment of RPE-1 cells, and a decrease in cells in the G0/G1 and S phases. After treatment, 37.4% of the rate cells were in the G2/M phase, compared to 16.3% in the control group. After 24h of H_2_O_2_ treatment, MSC-CM reversed the increased proportion of cells in the G2/M phase compared to basal group (27.2%). When the cells were treated with ADMSC-CM and LMSC-CM, the proportions of RPE-1 cells in the G2/M phase were reduced to 18.3% and 17.0%, respectively (Fig. 3).

**Figure 3.**
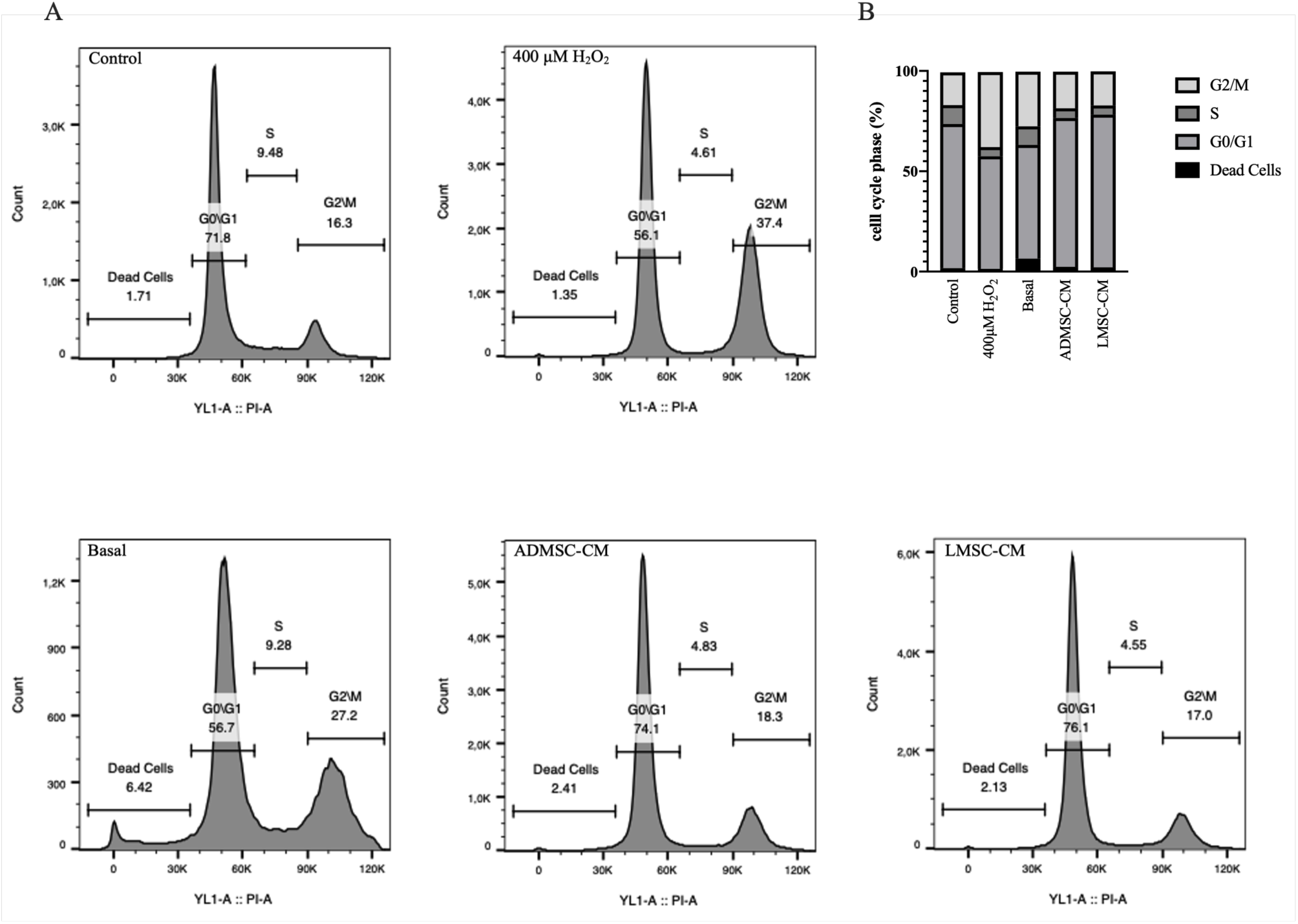
MSC-CM attenuates the G2/M phase arrest induced by oxidative stress. The cell cycle analysis of RPE-1 cells was determined by flow cytometry. The flow plots (A) and the quantification results as a graph (B) were shown.

### MSC-CM protected RPE cells by inhibiting p38 signaling

Senescence-associated secretory phenotype (SASP) refers to the tendency of cells stalled in the G2/M phase to release inflammatory cytokines in an attempt to attract immune cells (Giuliani et al., 2025; Grun et al., 2023). Thus, we decided to determine whether the regulation of cell cycle arrest also led to changes in the inflammatory response. The gene expression levels of IL-1β, IL-6, and IL-10 was determined using RT-qPCR. We observed that H_2_O_2_ treatment induced an increase in the gene expression levels of IL-1β, IL-6, and IL-10. Specifically, while the transcription of IL-1β continued to increase in the basal group, the gene expression levels of IL-6 and IL-10 exhibited a decreasing trend compared to the H_2_O_2_ group yet remained elevated relative to the control group. Treatment with ADMSC-CM resulted in a reduction in gene expression levels compared to the basal group, with IL-6 and IL-10 gene expression levels approaching those of the control group. Similarly, LMSC-CM treatment led to a decrease in IL-1β and IL-6 gene expression levels compared to both the basal and H_2_O_2_ groups, but IL-10 gene expression showed a significant increase compared to the basal and ADMSC-CM group (Fig. 4A). Next, we wanted to establish the ratio of p-p38 to p38, given that the majority of research showed that p38 MAPK regulates cytokine gene expression (Nahirnyj et al., 2013; Neuder et al., 2009). Increased phosphorylation levels of p38-MAPK in the H_2_O_2_ and basal group was significantly reversed with MSC-CM treatment (Fig. 4B).

**Figure 4.**
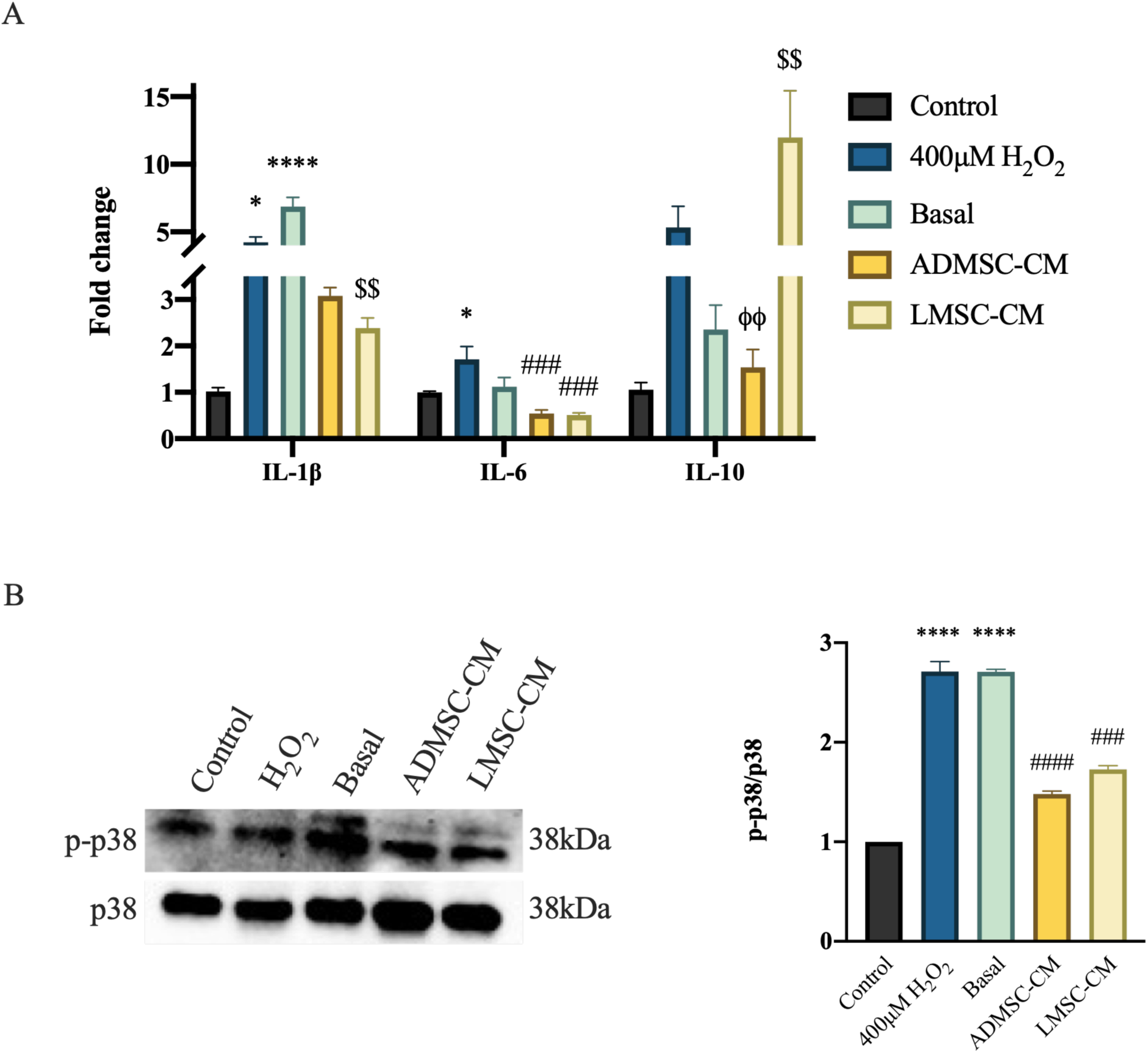
MSC-CM suppressed H₂O₂-induced inflammation. A) RT-qPCR results show the expressions of IL-1β, IL-6 and IL-10 genes. B) Western blot analysis for protein levels associated with the p38-MAPK pathway. Vinculin was used as an internal reference. Data shown are representative of 3 separate experiments and values are given as mean ± SEM. Statistical analysis was performed by one-way ANOVA and Kruskal-Wallis test. * compared to control group, # compared to H₂O₂ group, $ compared to basal group, φ compared to LMSC-CM group; for all comparisons * p<0.05, ** p<0.01, *** p<0.001, **** p<0.0001.

### MSC-CM Inhibited Apoptosis

The BCL2 family plays a crucial role in controlling cell survival (Hoang et al., 2022; Yang et al., 2019), and the activation of the p38 pathway is one of the mechanisms that precedes caspase activation and initiates apoptosis (Panuthai et al., 2023). This step represents the ultimate stage in stress signaling prior to caspase activation, which leads to cell death through apoptosis (Whitaker & Cook, 2021). Western blotting assays showed how apoptosis-associated proteins were impacted. We examined the expression of pro-apoptotic protein (Bax) and anti-apoptotic protein (Bcl-2) in RPE-1 cells. The Bax/Bcl-2 ratio significantly increased after H_2_O_2_ treatment, exhibiting a higher trend compared to the control in the basal group. However, with MSC-CM treatment, this ratio showed a similar level to the control group. Furthermore, the decrease observed with ADMSC-CM was significantly lower compared to the basal group (Fig. 5A).

**Figure 5.**
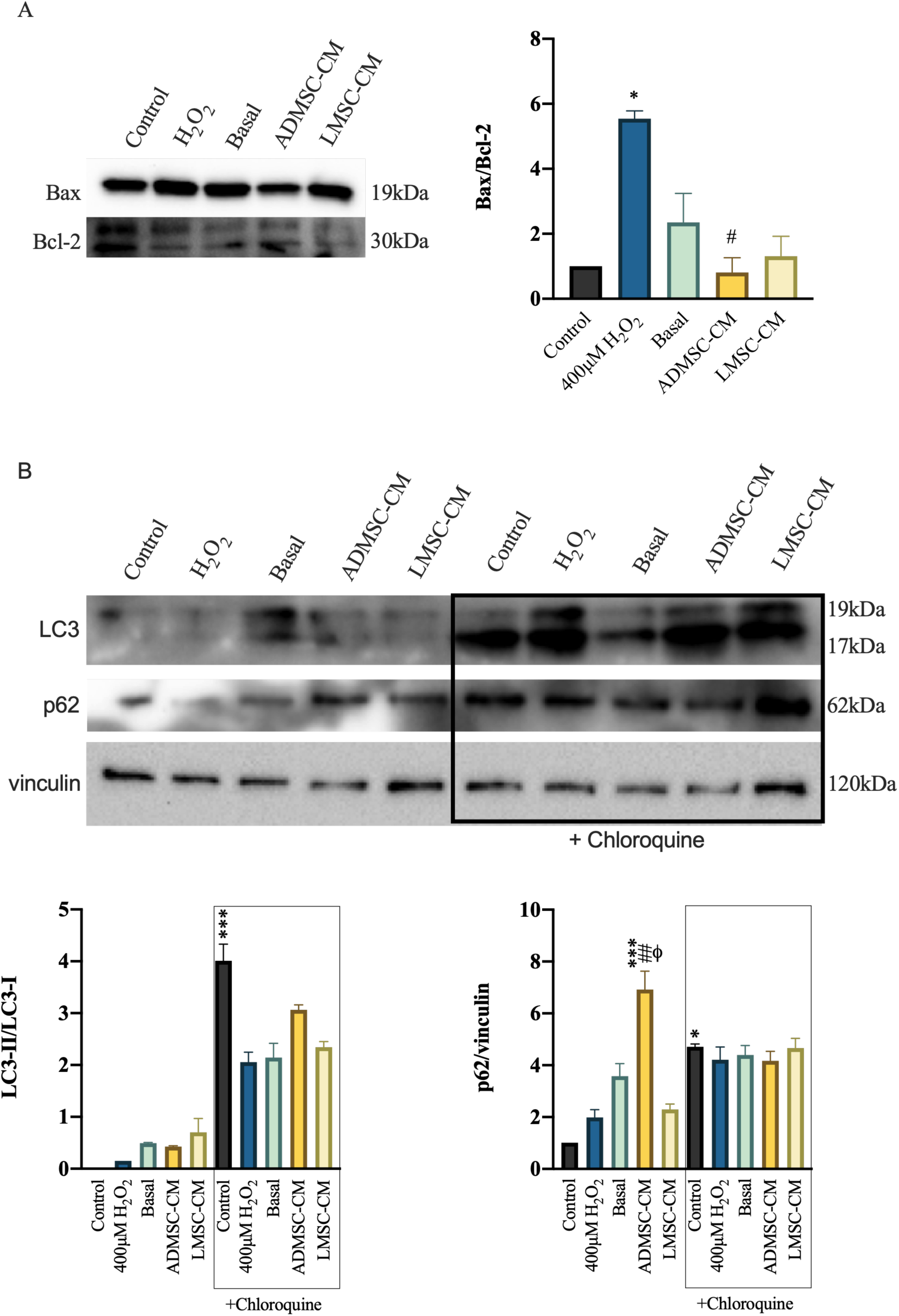
MSC-CM regulated apoptosis-associated proteins H₂O₂-treated cells. A) Western blot analysis of Bax and Bcl-2 were shown. Same blot was used to probe. The ratio of the Bax/Bcl-2 as determined by densitometry is illustrated in the graph. B) Analysis of autophagy dynamics after MSC-CM application in H₂O₂-treated RPE cells. LC3 and p62 immunoblots are shown. Vinculin is used as a normalization control. The ratios of the LC3-II/LC3-I and p62 levels as determined by densitometry are illustrated in the graphs. Data shown are representative of 3 separate experiments and values are given as mean ± SD. Statistical analysis was performed by Kruskal-Wallis test. * compared to control group, # compared to H₂O₂ group and, φ compared to LMSC-CM group; for all comparisons * p<0.05, ** p<0.01, *** p<0.001.

In order to determine if autophagy is involved in the protective effects of MSC-CM following H_2_O_2_ treatment, we analyzed the expression profiles of LC3 and p62 in RPE-1 cells treated with 400 µM H_2_O_2_. This analysis was conducted both with and without the presence of a well-known autophagy inhibitor, chloroquine. Increased autophagy activation is indicated by the conversion of LC3-I to LC3-II, whereas normal autophagic flux is indicated by the destruction of p62 at the end of the autophagy process. Western blot analysis identified the LC3II/I ratio, which increased with H_2_O_2_ treatment, continued to rise at the basal level; however, MSC-CM treatment did not show any difference compared to the basal level. ADMSC-CM treatment resulted in a significant increase in p62 levels compared to the control, H_2_O_2_, and LMSC-CM groups. Upon chloroquine administration, the control group exhibited a significant increase in both the LC3II/LC3I ratio and p62 levels compared to the non-chloroquine condition. In contrast, ADMSC-CM treatment showed levels closer to the control group than to the H_2_O_2_, basal, and LMSC-CM groups. No differences in p62 levels were observed between the groups (Fig. 5B).

### MSC-CM did not show any pro-angiogenic effect

Proangiogenic substances, such as VEGF and vasculogenic and inflammatory cytokines, are secreted into the choroidal space by the RPE in cases of AMD. This might initiate angiogenesis, which would result in choroidal neovascularization (CNV) (Vega et al., 2021).

We used ELISA to assess the amount of VEGF-A in the culture media of RPE-1 treatments in order to observe whether MSC-CM treatment demonstrates pro-angiogenic effects. There were no noticeable differences in the secreted VEGF-A levels according to the ELISA data (Fig. 6A). Then, RPE-1 culture media was used as HUVEC treatment medium. We used the RPE-1 treatment media combined in a 1:1 ratio with HUVEC medium for this purpose. The HUVEC medium was combined 1:1 with DMEM/F12 medium to form the “Only medium” group. MTT assay showed that the culture media from the ADMSC-CM and LMSC-CM groups significantly reduced HUVEC vitality in contrast to the 400 µM H_2_O_2_ group (Fig. 6B). We also evaluated how culture media affected the HUVECs’ ability for migration and invasion. The H_2_O_2_ and basal groups in the migration assay showed closure, whereas the MSC-CM groups showed closure rates that were close to those of the control groups (Fig. 6C). MSC-CM groups showed a statistically significant decrease in wound closure compared to the basal group (Fig. 6E). Then, the invasion ability of the cells was examined by Transwell assay. After 6 hours of using RPE-1 culture medium as chemoattractant medium, the cells were fixed and stained with crystal violet and photographed (Fig. 6D). The results of the analysis showed increased cell invasion in the H_2_O_2_ and basal groups. However, MSC-CM treatment reversed this increase (Fig. 6E). This experiment shows that there are no pro-angiogenic effects from MSC-CM treatment.

**Figure 6.**
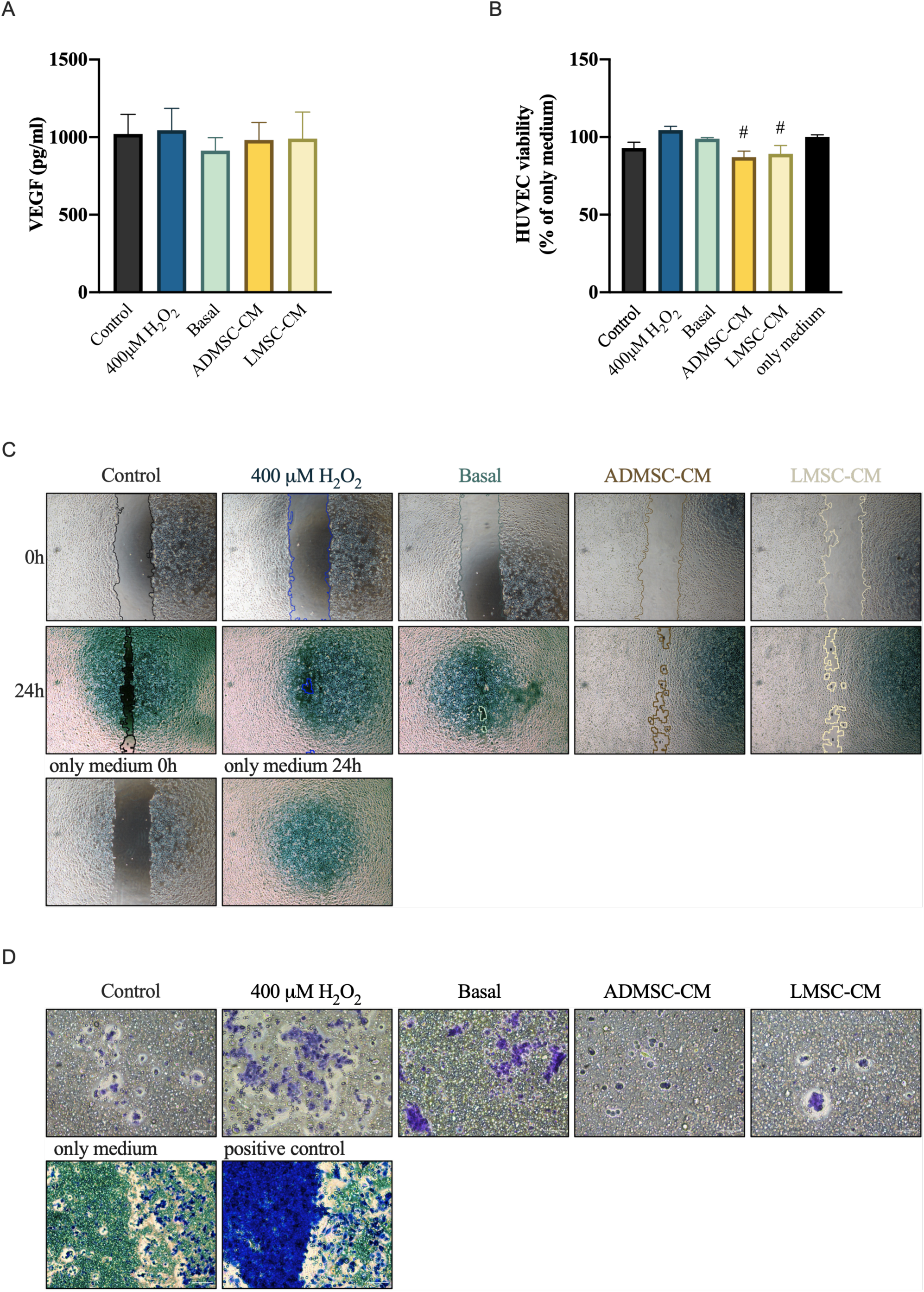
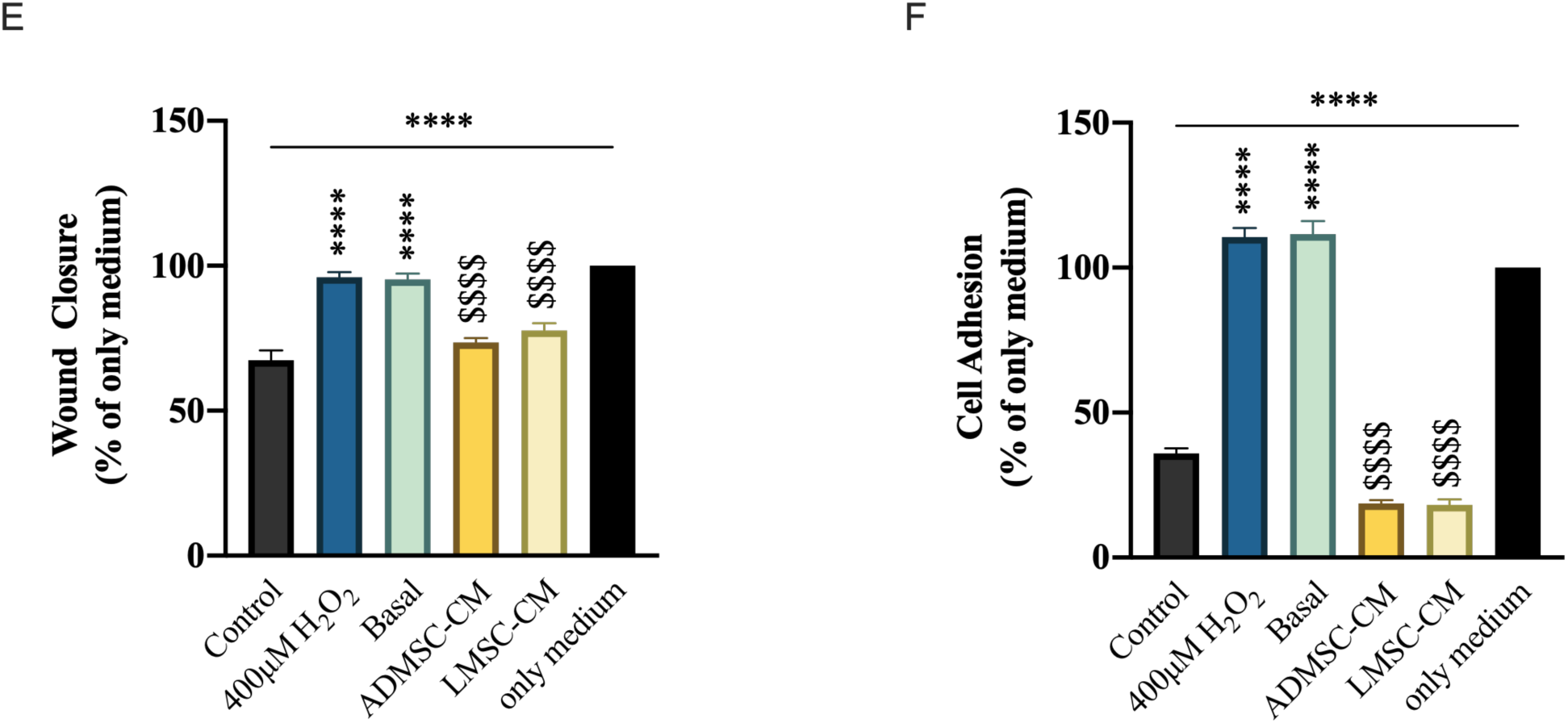
MSC-CM treatment did not increase VEGF-A secretion and significantly reduced HUVEC viability, migration, and invasion. A) ELISA analysis of VEGF-A production in RPE-1 culture media. B) MTT analysis of cell viability in HUVECs after 24h application of RPE-1 culture media. The results are expressed as percentage of only medium. C, E) Representative photomicrographs and analysis of HUVEC wound closure by scratch assay. D, F) Representative photomicrographs and analysis of HUVEC invasion by Transwell assay. As a positive control, medium containing 10% FBS was used. Data shown are representative of 3 separate experiments and values are given as mean ± SEM. Statistical analysis was performed by One-way ANOVA. * compared to control group, # compared to H₂O₂ group and, $ compared to basal group; for all comparisons * p<0.05, **** p<0.0001.

## DISCUSSION

The homeostatic regulation of the retinal pigment epithelium (RPE) is a complex process, and its dysfunction resulting from persistent oxidative stress is a main cause of retinal degeneration (Tsang & Sharma, 2025; Xu et al., 2023). In this study, we provide a comparative analysis of the secretomes from Adipose-derived (ADMSC) and Limbal-derived (LMSC) Mesenchymal Stem Cells, evaluating their efficacy in rescuing RPE-1 cells from H_2_O_2_-induced dysfunction. Our choice of LMSCs as a comparator to the clinically established ADMSCs was predicated on the ‘niche-specific’ hypothesis: that cells residing within the ocular microenvironment possess a proteomic specifically evolved to handle the high oxidative demands of the eye (Altug et al., 2024; Shukla et al., 2020; Tomasello et al., 2025)

Our results demonstrate that both ADMSC-CM and LMSC-CM significantly enhance RPE-1 viability following an oxidative insult, consistent with the established protective role of the MSC secretome in ocular models (Liu et al., 2025; Suthumchai et al., 2025; Yu et al., 2024). Even so, the therapeutic narrative is not fully captured by the survival data alone. Interestingly, while cell survival increased, Ki67 staining showed no significant shift in proliferation. This is a vital safety hallmark; in the subretinal space, uncontrolled proliferation can lead to deleterious complications such as subretinal fibrosis or epiretinal membrane formation (Lee et al., 2016; Vannay et al., 2005). Rather, it seems that the MSC secretome gives priority to the restoration of existing RPE cells’ metabolism and structure (Tang et al., 2023). This strategy guarantees the preservation of the blood-retinal barrier without exacerbating the fibrovascular issues that are frequently observed in advanced AMD.

The progression of RPE-1 cells into a G2/M phase arrest subsequent to DNA damage is a well-documented survival strategy (Nikolakopoulou et al., 2021), yet it is a double-edged sword. If this arrest is prolonged, cells enter senescence and adopt a senescence-associated secretory phenotype (SASP), characterized by the chronic release of pro-inflammatory cytokines like IL−1β and IL−6 (Blasiak, 2020; Lazzarini et al., 2022; Ou et al., 2024). Our data demonstrate that both MSC secretomes, especially LMSC-CM, significantly alleviate this arrest. This reversal indicates a dynamic control of the G2/M checkpoint, perhaps via the regulation of CDK inhibitors like p21 and p27 (Do et al., 2019; Li et al., 2011). By inhibiting the progression to a persistent senescent state, MSC-CM therapies significantly reduce the risk of chronic, secondary inflammation in the retinal milieu. This molecular rescue is further evidenced by the suppression of the p38 MAPK pathway. One of the essential components of cell cycle progression in retinal pathologies is p38 MAPK, and its inhibition results in cell cycle arrest (Chen et al., 2018; Yamaguchi et al., 2002). Additionally, a phosphorylation cascade in the p38 pathway is started by the overexpression of stress-response genes such Gadd45g, which acts as a crucial junction that results in either repair or apoptosis (Muraleva & Kolosova, 2023; Zhu et al., 2009). The significant decrease in p38 phosphorylation after CM therapy establishes a direct mechanistic connection to the stabilization of the Bax/ Bcl−2 ratio. By attenuating this stress-activated protein kinase signaling, the secretome effectively protects the cell from reaching the’point of no return’ in the apoptotic cascade (Whitaker & Cook, 2021).

A significant aspect of our findings is the differential regulation of the autophagic protein p62 (SQSTM1). Autophagy is an essential cellular recycling mechanism in the retinal pigment epithelium, and its dysregulation is directly associated with the buildup of lipofuscin and the subsequent advancement of age-related macular degeneration (AMD) (Kaarniranta et al., 2023). We assessed the autophagic response to figure out if the protective impact of the secretome was facilitated by the clearance of oxidative debris. Our results showed that while H_2_O_2_ increased the LC3II/I ratio ADMSC-CM treatment uniquely led to a significant accumulation of p62.

Typically, p62 accumulation is interpreted as a blockade in autophagic flux (Bjørkøy et al., 2009), however, in the context of acute oxidative stress, p62 serves a more complex, non-canonical role.It functions as an essential adaptor protein that stabilizes Nrf2 (Nuclear Factor Erythroid 2-related factor 2) by sequestering its negative regulator, Keap1 (Al Saihati et al., 2024; Ichimura et al., 2013).We suggest that the elevation of p62 seen with ADMSC-CM therapy may indicate the strategic transition towards reinforcing the cell’s intrinsic antioxidant defenses. The secretome likely promotes Nrf2 nuclear translocation, hence enhancing the transcription of antioxidant response elements (ARE) (Liu et al., 2022), providing a layer of protection that complements the direct dampening of p38 MAPK signaling (Yang et al., 2022). This indicates a divergence in the therapeutic approaches of the two MSC sources: whereas LMSCs are very successful in alleviating cell cycle arrest and inflammation, ADMSCs may provide enhanced protection by activating the RPE’s endogenous antioxidant system via the p62-Nrf2 pathway. The combined strategy of LMSC’s regeneration signals and ADMSC’s metabolic support highlights the possibility for a synergistic cell-free therapy in treating retinal degeneration.

The clinical development of AMD typically evolves from the dry atrophic form to wet neovascular AMD, characterized by the uncontrolled growth of leaky vessels from the choriocapillaris into the sub-retinal space (Handa, 2012). Consequently, an optimal treatment drug must not only rescue RPE cells but also stabilize the underlying vascular milieu. Our HUVEC migration experiments demonstrated that both ADMSC-CM and LMSC-CM have notable anti-angiogenic characteristics, greatly restricting endothelial cell motility. This action which is protecting the RPE against oxidative death while simultaneously suppressing the pro-angiogenic signals that drive choroidal neovascularization (CNV) is a major therapeutic advantage.We hypothesize that this impact is facilitated by the regulated release of anti-angiogenic substances such as Pigment Epithelium-Derived Factor (PEDF) and certain microRNAs contained in MSC-derived exosomes (Mead et al., 2020; Sheibani et al., 2019; Toh et al., 2018). By modulating the extracellular milieu, MSC-CM acts as a biologic stabilizer, potentially slowing the devastating transition to the neovascular stages of retinal disease. The integrated summary of proposed pathways is illustrated in Figure 7.

**Figure 7.**
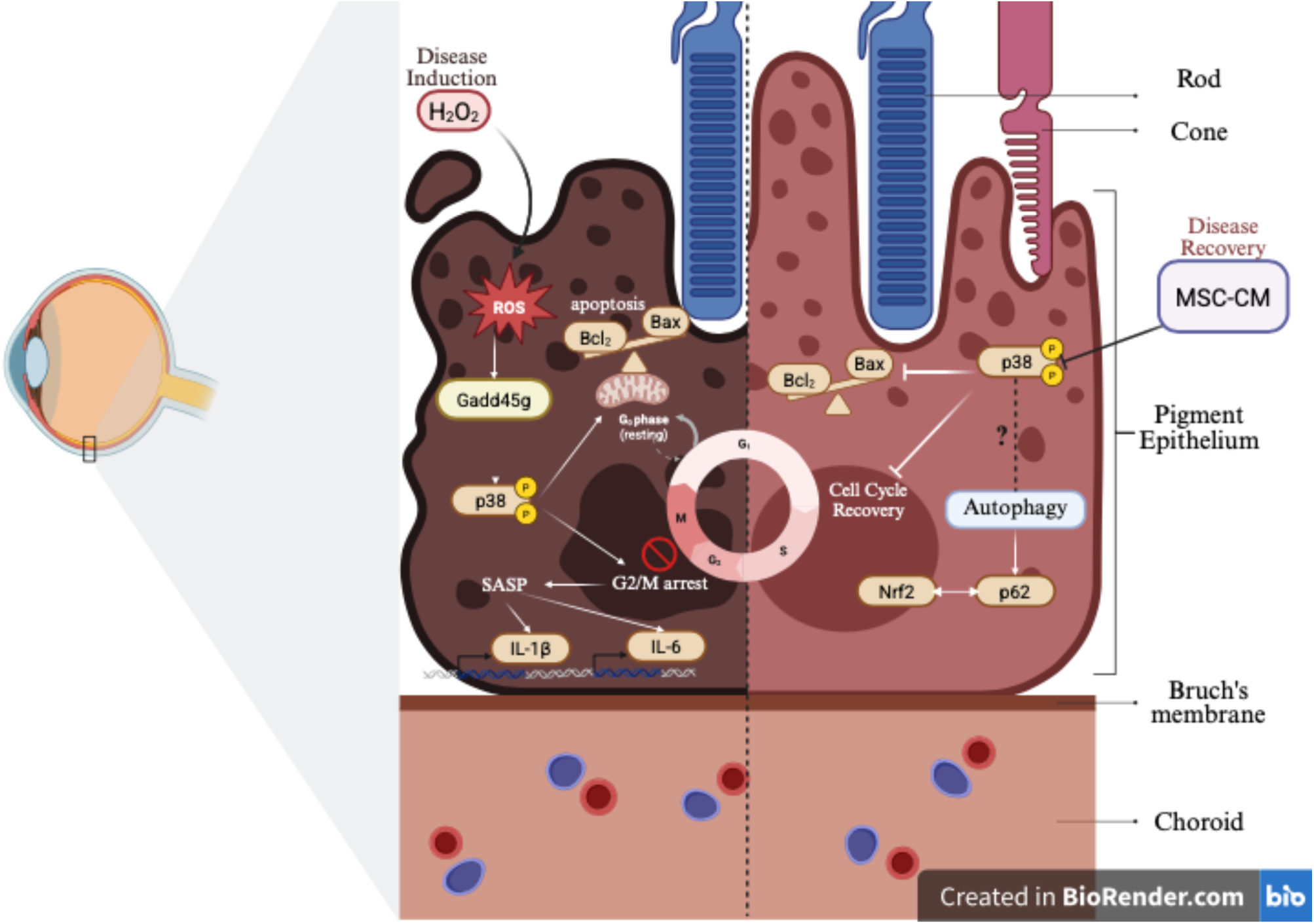
Proposed Molecular Mechanism of MSC-CM Mediated RPE Rescue. Integrated schematic representation of the therapeutic effects of ADMSC and LMSC secretomes on oxidative-stressed RPE-1 cells. Left Panel (Disease State): Exposure to H₂O₂ triggers an acute stress response characterized by the activation of the Gadd45g/p38 MAPK signaling cascade. This activation facilitates a shift in the Bax/Bcl−2 ratio towards a pro-apoptotic state and induces a robust G2/M phase cell cycle arrest. Stalled cells in the G2/M phase contribute to the Senescence-Associated Secretory Phenotype (SASP), resulting in the release of pro-inflammatory cytokines such as IL-1β and IL-6, while simultaneously creating a pro-angiogenic environment that promotes HUVEC migration. Right Panel (Treatment Intervention): Treatment with MSC-conditioned media (MSC-CM) effectively intercepts these pathological pathways. The secretome suppresses p38 MAPK phosphorylation, thereby stabilizing mitochondrial integrity and preventing apoptosis. Crucially, MSC-CM facilitates the resolution of G2/M arrest, restoring homeostatic cell cycle progression. Notably, ADMSC-CM uniquely upregulates p62 (SQSTM1), potentially priming the Nrf2-mediated antioxidant defense to neutralize ROS. Furthermore, the secretome exerts a potent anti-angiogenic effect by inhibiting endothelial cell motility, collectively contributing to the stabilization of the retinal and vascular microenvironment (Created in BioRender.com).

We acknowledge that this in vitro model, while mechanistically informative, lacks the three-dimensional complexity of the blood-retinal barrier and the interplay with the choroidal vasculature. However, these findings establish a strong foundation for the next logical step in clinical translation. The instability of liquid-form secretomes remains a hurdle for widespread use; therefore, our upcoming research is dedicated to evaluating the bioactivity of lyophilized (freeze-dried) ADMSC-CM. Liyophilization not only ensures the long-term stability of growth factors and exosomal microRNAs but also allows for the development of ‘off-the-shelf’ regenerative therapies. Our future goals include characterizing the proteomic stability of these freeze-dried products and testing their efficacy in in vivo models to determine if this cell-free approach can successfully prevent vision loss in a complex biological system.

## ACKNOWLEDGEMENTS

The authors gratefully acknowledge use of the services and facilities of the Koç University Research Center for Translational Medicine (KUTTAM), funded by the Presidency of Strategy and Budget. The content is solely the responsibility of the authors and does not necessarily represent the official views of the Presidency of Strategy and Budget.

